# Continuous amyloid staging complements global amyloid burden in early Alzheimer’s disease

**DOI:** 10.64898/2026.10.03.756367

**Authors:** Haoyuan Liu, Mingzhao Tong, Shiaofen Fang, Elizabeth C. Mormino, Reisa A. Sperling, Paul Aisen, Rachel F. Buckley, Paul K. Crane, Timothy J. Hohman, Andrew J. Saykin, Jingwen Yan, the Alzheimer’s Disease Neuroimaging Initiative

## Abstract

**Introduction:** Early Alzheimer’s disease (AD) staging remains challenging because clinical categories and global amyloid PET measures provide only coarse characterization of disease progression. We developed a continuous amyloid staging score from regional amyloid PET to complement global amyloid burden.

**Methods:** Using longitudinal florbetapir PET from the Alzheimer’s Disease Neuroimaging Initiative (ADNI), we trained an end-to-end deep learning model to directly estimate a continuous amyloid staging score. We evaluated longitudinal consistency, its relationship with global amyloid burden, regional contributors to early-stage variation, and whether the staging score stratified subsequent memory decline among participants matched for baseline global amyloid burden. Cognitive analyses were replicated in A4+LEARN.

**Results:** The staging score better preserved longitudinal visit order than existing amyloid staging approaches. During early disease stages, it captured regional amyloid variation despite relatively stable global amyloid burden. Among participants with similar baseline global amyloid burden, higher staging scores were associated with faster subsequent memory decline in both ADNI and A4+LEARN.

**Discussion:** A continuous amyloid staging score complements global amyloid burden for early AD staging and cognitive risk stratification.

## 1. Introduction

Alzheimer’s disease (AD) develops gradually over many years before the onset of overt cognitive impairment, creating a critical window for early detection and intervention ^1^. Amyloid-β accumulation is one of the earliest pathological changes in AD and can be detected in vivo using amyloid positron emission tomography (PET), providing a biological marker of preclinical disease ^2^. The Centiloid scale provides a standardized continuous measure of global cortical amyloid burden ^3^. Although widely used in clinical trials, it summarizes regional amyloid PET measurements into a single global value. During the earliest stages of disease, amyloid accumulation is subtle and initially limited to a small number of brain regions. Consequently, averaging across the cortex may dilute these localized changes before they substantially influence the global measure, potentially limiting sensitivity to early disease progression.

Disease progression models have been developed to leverage regional amyloid PET measurements for more refined disease staging. Event-based models (EBMs) infer the temporal ordering of abnormalities across a predefined set of regional biomarkers, providing refined but still discrete disease stages ^4–6^. More recently, continuous trajectory-learning methods, originally developed in single-cell biology to reconstruct cellular differentiation trajectories ^7,8^, have been adapted to estimate disease progression from molecular and neuroimaging biomarkers in AD ^9^. SLOPE further incorporated longitudinal follow-up information into this framework by introducing an ordering constraint during representation learning, enabling continuous amyloid staging of previously unseen individuals while proposing quantitative metrics to evaluate staging robustness ^10^. Compared with global amyloid burden, SLOPE demonstrated improved sensitivity to subtle regional amyloid changes during the early stages of disease. However, the final staging score is obtained through multiple sequential processing steps rather than being learned directly. As a result, preservation of longitudinal ordering is enforced during an intermediate processing step rather than during estimation of the final staging score, potentially limiting temporal consistency across longitudinal follow-up visits.

In this study, we developed DAPS (Direct Amyloid PET Staging), an end-to-end deep learning framework that directly estimates a continuous amyloid staging score from regional amyloid PET. By directly optimizing the final staging score to preserve longitudinal ordering, DAPS eliminates the need for downstream manifold embedding and trajectory fitting. We hypothesized that DAPS would improve temporal consistency of longitudinal staging while providing complementary information to global amyloid burden for early disease staging. We further evaluated whether the DAPS staging score stratifies subsequent memory decline among individuals with similar baseline global amyloid burden and whether these findings generalize to the independent A4+LEARN cohort.

## 2. Methods

### 2.1 Study cohorts and PET features

All analyses were performed using harmonized participant-level variables and amyloid PET imaging data provided by the Alzheimer’s Disease Sequencing Project (ADSP) Phenotype Harmonization Consortium ^11,12^. The primary cohort comprised longitudinal florbetapir amyloid PET data from the Alzheimer’s Disease Neuroimaging Initiative (ADNI), a multicenter observational study of Alzheimer’s disease biomarkers and clinical progression ^13^. The analytic dataset included 1,161 participants after restricting analyses to those with complete cortical regional amyloid measurements, and the non-Hispanic White subgroup shared with the validation cohort. Baseline diagnoses and longitudinal follow-up are summarized in Figure 1.

**Figure 1.**
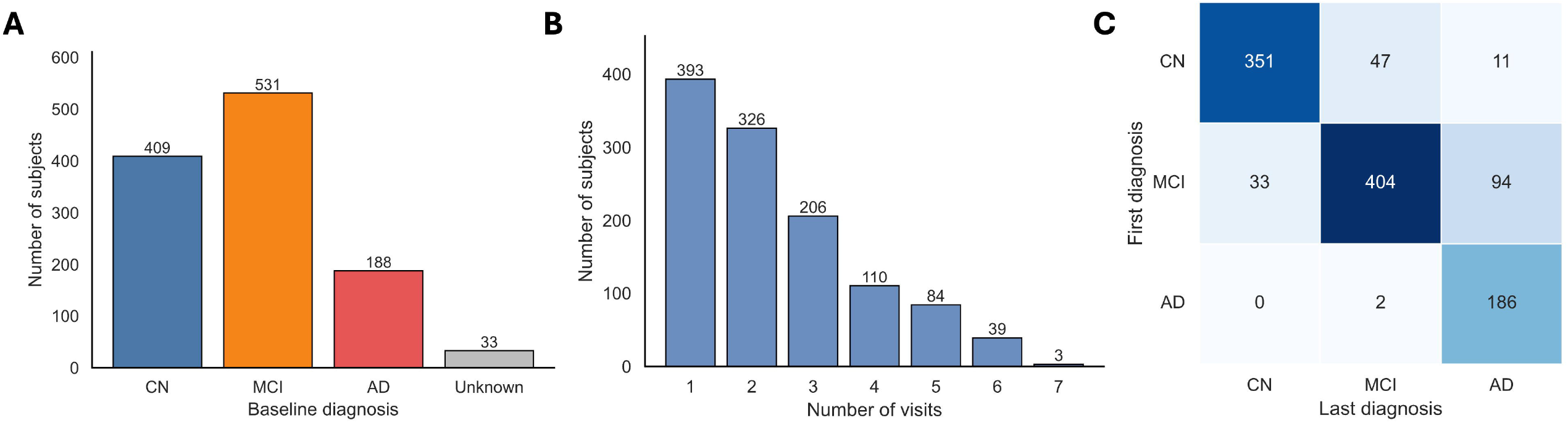
Overview of the ADNI cohort. (A) Baseline diagnostic distribution. (B) Distribution of amyloid PET visits per participant. (C) Diagnostic transitions between the first and last visits among participants with longitudinal follow-up.

External validation was performed using harmonized PET data from the A4 and LEARN cohorts ^14,15^ after applying the same demographic restriction (non-Hispanic White subgroup). To avoid potential treatment effects on amyloid progression and cognition, only placebo participants from the A4 trial were included, totaling 3,414 participants in the validation cohort. Detailed demographic information of the study cohorts is provided in Table 1.

In both cohorts, standardized uptake value ratios (SUVRs) from 68 cortical regions were analyzed for each PET scan. Regional SUVRs were adjusted for age, sex, and education using regression coefficients estimated from baseline cognitively normal participants in the training cohort. The same adjustment was then applied unchanged to the independent test cohort and the A4+LEARN cohort. In ADNI, participants with longitudinal PET were randomly divided at the subject level into training (n = 777) and independent test (n = 384) cohorts, while all single-visit participants were included in the training cohort.

In the downstream cognitive decline analysis, longitudinal cognition was assessed using the ADSP harmonized memory composite (PHC_MEM) (ADNI and A4+LEARN) and the Preclinical Alzheimer Cognitive Composite (PACC) (A4+LEARN only), which is known for greater sensitivity of subtle cognitive decline in cognitively unimpaired individuals.

### 2.2 Direct Amyloid PET Staging (DAPS)

DAPS is an end-to-end deep learning framework that directly estimates a continuous amyloid staging score from regional amyloid PET measurements. The framework consists of an encoder that compresses regional amyloid PET measurements into a one-dimensional staging score and a decoder that reconstructs the original regional amyloid PET pattern. Reconstruction regularizes the learned representation, while longitudinal follow-up visits supervise the staging score through an ordering constraint.

Let *x*_*ik*_ ∈ ℝ^68^ denote the regional amyloid PET measurements for participant *i* at visit *k*. The encoder *f*_*θ*_ maps the input to a scalar latent staging score,

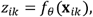

and the decoder *g* _*ϕ*_ reconstructs the regional amyloid PET measurements,

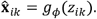

The latent staging score was transformed into a normalized staging score ranging from 0 to 1 using a sigmoid transformation followed by min-max normalization based on the training cohort,

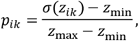

where *z*_min_ and *z*_max_ denote the minimum and maximum transformed latent scores in the training cohort. The same transformation was subsequently applied to the independent ADNI test cohort and the external A4+LEARN cohort. The normalized staging score *p*_*ik*_ was used throughout subsequent analyses.

To preserve longitudinal disease progression, DAPS incorporates a longitudinal ordering loss that directly constrains the final staging score. For participants with multiple PET visits ordered by acquisition age, staging scores are encouraged to remain stable or increase across successive visits. Specifically,

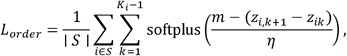

where *S* denotes participants with longitudinal follow-up, is the positive ranking margin, and *η* controls the smoothness of the penalty. This differentiable objective penalizes violations of longitudinal ordering while allowing small visit-to-visit fluctuations.

The overall training objective combines reconstruction loss, longitudinal ordering loss, and latent-scale regularization,

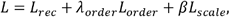

where

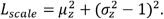

Here, *L*_*rec*_ is the mean squared reconstruction loss, and *μ*_*z*_ and 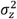 denote the batch mean and variance of the latent staging scores. The scale regularization encourages the latent scores to remain centered around zero with unit variance, stabilizing optimization and maintaining a consistent scale for the longitudinal ordering constraint. Model training was fully self-supervised without using diagnostic labels. The final DAPS model consisted of a fully connected encoder-decoder architecture with a one-dimensional latent staging score. Specifically, the encoder used layer dimensions [68, 512, 128, 1], and the decoder used the mirrored architecture [1, 128, 512, 68]. ReLU activation was applied after each hidden linear layer, with no dropout. Linear-layer weights were initialized orthogonally and biases were initialized to zero. The model was trained for 300 epochs using AdamW with a learning rate of 3e-5, weight decay of 1e-5, and batch size of 256. The training objective combined mean squared reconstruction loss, a longitudinal order loss weighted by 20, and a latent scale regularization term weighted by 0.5. The order loss used a fixed margin of 0.2 and a softplus temperature of 0.1. After training, the scalar encoder output was transformed by a sigmoid function and min-max scaled using the training-set range to obtain the final staging score.

### 2.3 Model training and evaluation

Participants with longitudinal PET were randomly divided at the subject level into training (n = 777) and independent test (n = 384) cohorts, while all single-visit participants were included in the training cohort. Subject-level partitioning prevented data leakage between training and testing. Model architecture and hyperparameters were determined using the training cohort only (Supplementary Methods). The finalized DAPS model was subsequently evaluated in the independent ADNI test cohort and directly applied to the external A4+LEARN cohort without retraining. DAPS was compared with SLOPE ^10^, and also simple autoencoder (AE-UMAP) without longitudinal ordering constraint while preserving the same downstream staging workflow. Global amyloid burden (Centiloid) was normalized to the same [0,1] range for comparison.

Temporal consistency was evaluated in the independent ADNI test cohort. A violation occurred when the staging score decreased between two consecutive PET visits. The violation ratio quantified the proportion of ordering violations, while the violation gap measured the magnitude of these violations. Lower values indicate better preservation of longitudinal ordering.

### 2.4 Association of DAPS staging score with Centiloid

We fit a continuous segmented regression relating Centiloids to the normalized staging score. The breakpoint was estimated in the ADNI training set and the estimated breakpoint was at the DAPS staging score of 0.370. The same breakpoint was then applied to the ADNI test set and A4+LEARN cohort, and was used to define the early and late staging intervals for subsequent analyses.

To identify regional features contributing to early DAPS staging variation, we used the trained decoder to trace reconstructed regional PET features across the early pseudotime interval from 0 to the training-set breakpoint 0.37. For each cortical feature, the contributing score was defined as the mean decoder gradient over 200 evenly spaced pseudotime values in this interval. Regions with larger positive scores therefore showed stronger amyloid increases during early progression along the estimated progression trajectory.

### 2.5 Longitudinal cognitive decline analyses

The downstream analyses evaluated whether the baseline DAPS staging score could stratify future cognitive decline beyond global amyloid burden. Analyses were restricted to participants with baseline CN or MCI diagnosis and baseline Centiloids ≤60, approximately corresponding to the upper Centiloid limit at the training-derived breakpoint, before which DAPS is expected to provide complementary staging information beyond global amyloid burden. Participants were classified into early- and late-stage groups using the training-derived staging breakpoint (0.370). To isolate the contribution of the DAPS staging score beyond global amyloid burden, participants were matched on baseline Centiloids using ten pooled quantile bins, with equal numbers retained from the early- and late-stage groups within each bin to ensure comparable baseline Centiloid distributions. After matching participants on baseline Centiloids and adjusting for practice effects following ^16^, longitudinal cognitive trajectories were evaluated using the following linear mixed-effects model,

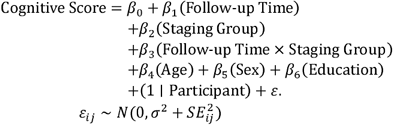

where Staging Group indicates the early- or late-stage group defined by the training-derived breakpoint. (1 | Participant) represents a participant-specific random intercept. The observation-specific measurement precision of each psychometrically derived PHC_MEM score was incorporated into the model using its estimated standard error (SE)^17^. For PACC score in the A4+LEARN cohort, measurement precision (SE) was not provided and was set to zero. The residual term *ε* has variance equal to the sum of the estimated measurement variance and an additional residual variance. The interaction between follow-up time and staging group was the primary test of whether CN/MCI participants in the late-stage group exhibited faster cognitive decline than those in the early-stage group. The same matching strategy and statistical framework were applied to both the ADNI and A4+LEARN cohorts.

## 3. Results

### 3.1 DAPS staging score improves temporal consistency of longitudinal amyloid staging

The DAPS staging score increased progressively across diagnostic groups in the independent ADNI test cohort (Figure 2A). Because diagnostic labels were not used during model training, this pattern indicates that our self-supervised framework learned a biologically meaningful staging representation consistent with the clinical progression of Alzheimer’s disease. Temporal consistency was then evaluated by examining whether the staging score preserved the observed temporal order of longitudinal PET visits. DAPS achieved the lowest violation ratio (0.3424), outperforming both SLOPE (0.4475) and the simple autoencoder (AE-UMAP, 0.4561) (Figure 2B), indicating better preservation of longitudinal visit order. In addition, DAPS also produced the smallest violation gaps among all methods (Figure 2C), indicating that longitudinal ordering errors were not only less frequent but also smaller in magnitude. Together, these findings demonstrate that DAPS provides more temporally consistent longitudinal amyloid staging at both the population and individual levels.

**Figure 2.**
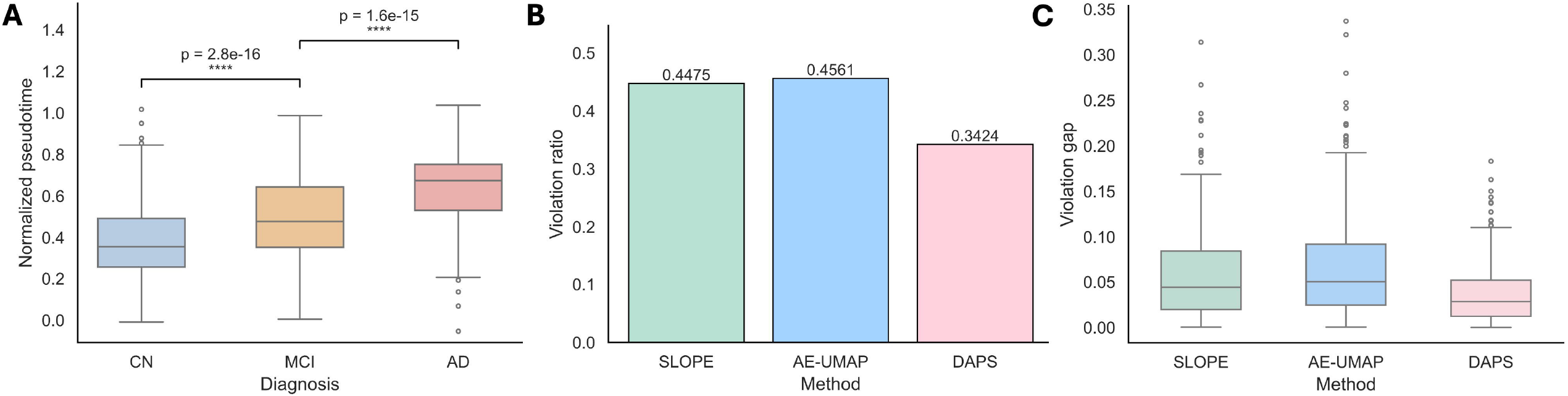
Temporal consistency of DAPS in the independent ADNI test cohort. (A) Distribution of the DAPS staging score across diagnostic groups. (B) Violation ratio of DAPS, SLOPE, and the simple autoencoder (AE-UMAP). (C) Violation gap distributions. Lower violation ratios and smaller violation gaps indicate better preservation of longitudinal visit order.

### 3.2 DAPS staging score complements global amyloid burden in early-stage AD

The relationship between the DAPS staging score and global amyloid burden was nonlinear across the ADNI training, ADNI test, and A4+LEARN cohorts (Figure 3A–C). A consistent transition point was observed, after which the staging score and Centiloids increased in parallel. In the ADNI training cohort, segmented linear regression identified a breakpoint at the staging score of 0.370. Before the breakpoint, the staging score showed little association with Centiloids (ADNI training: *R*^2^ = 0.004, *p* = 0.178), whereas a substantially stronger positive association was observed after the breakpoint (ADNI training: *R*^2^ = 0.44 *p* = 7.97 × 10^−136^). More specifically, during the early stage (DAPS score <0.37), the DAPS staging score continued to increase while global amyloid burden remained relatively stable. When the same breakpoint was applied to the held-out ADNI test set and independent A4+LEARN cohort, the transition from minimal to strong association was consistently observed, supporting the generalizability of the identified staging transition. These findings indicate that, during the early stage, DAPS may capture complementary information beyond global amyloid burden.

**Figure 3.**
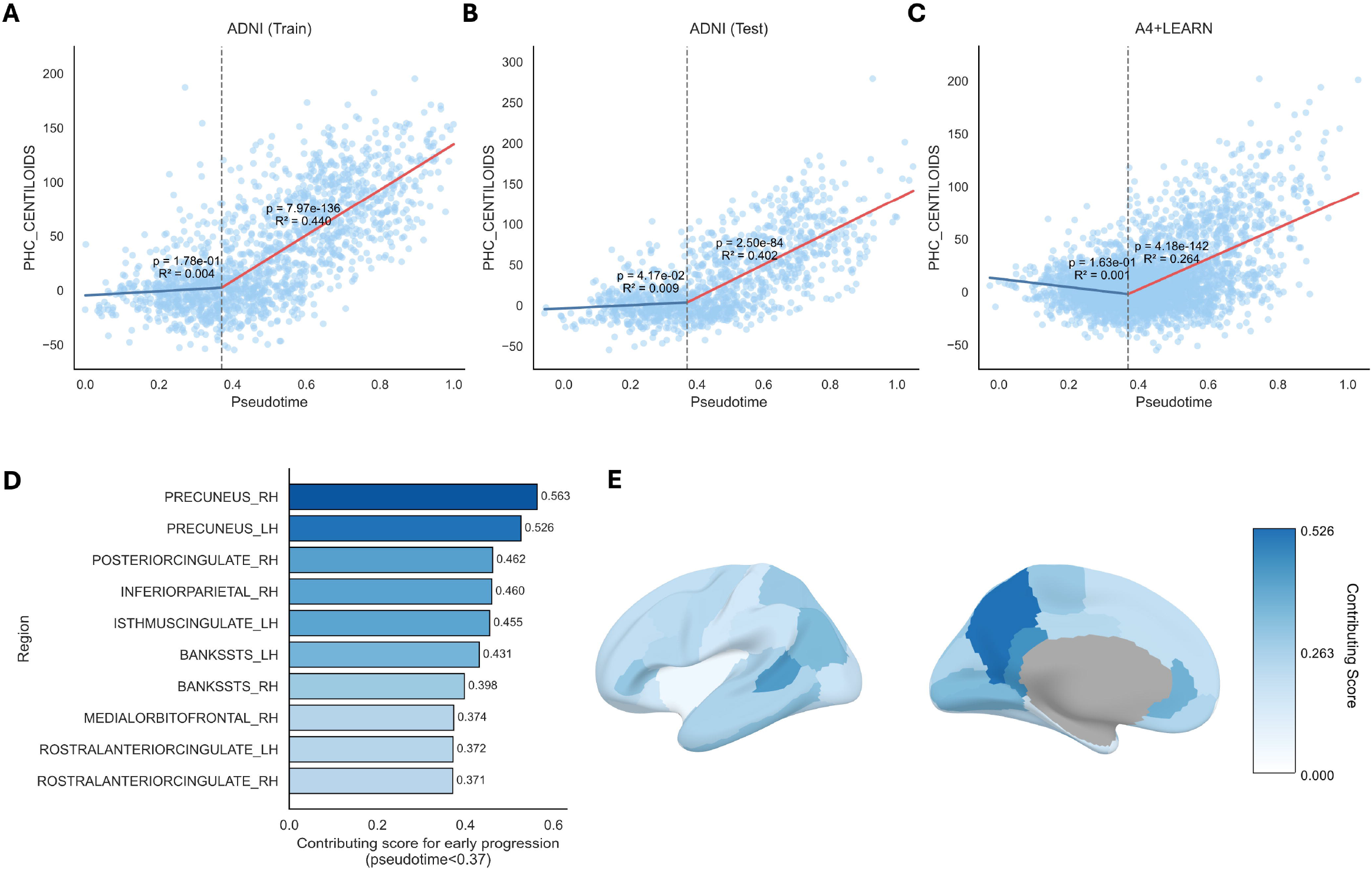
Relationship between the DAPS staging score and global amyloid burden. (A–C) Piecewise relationship between the DAPS staging score and Centiloids in the ADNI training, ADNI test, and A4+LEARN cohort. The dashed vertical line indicates the break point identified in the ADNI training set. (D) Ranked regional contribution scores during the early staging interval (before breakpoint 0.37). (E) Cortical map of top regions contributing to early DAPS staging score.

We further investigated the regional amyloid changes associated with the early staging interval (staging score < 0.370) using gradient analysis (Figure 3D–E). Each cortical region was assigned a contribution score that summarizes how strongly its regional amyloid burden changes with increasing staging score. Larger contribution scores therefore indicate regions whose amyloid accumulation changes drive the change of early DAPS staging score. We observed that not all cortical regions uniformly contributed to the early progression (Figure 3D–E), with higher values localized to a subset of cortical regions. The highest contribution scores were observed in the precuneus and posterior cingulate, both of which have been previously identified as early sites of amyloid accumulation over the course of Alzheimer development.

### 3.3 Late-stage participants showed faster memory decline despite matched Centiloids

To evaluate whether the baseline DAPS staging score provides complementary prognostic information beyond global amyloid burden, CN/MCI participants in ADNI test set and participants in the independent A4+LEARN cohort were divided into early- and late-stage groups using the staging breakpoint (0.370). The two groups were then matched in baseline Centiloids to ensure comparable baseline global amyloid burden. Matching produced nearly identical baseline Centiloids distributions in both ADNI and A4+LEARN (Figure 4A–B). Despite similar baseline global amyloid burden, late-stage declined significantly faster than early-stage participants in both cohorts. In ADNI test set, the late-stage group declined more rapidly on the harmonized memory score PHC_MEM than the early-stage group (*β* = 0.0299 PHC_MEM units/year, 95% CI [−0.0423], −0.0174, *p* = 2.56 × 10^−6^; Figure 4C). The same pattern held in A4+LEARN, where the late-stage group also showed faster decline on PHC_MEM (annualized *β* = −0.0234 PHC_MEM units/year, 95% CI [*β* = −0.0381, −0.0086, *p* = −0.0019; Figure 4D). After removing the measurement precision of PHC_MEM scores in the linear mixed model, stratification remained in both ADNI (*β* = −0.0304 PHC_MEM units/year, 95% CI [−0.0431, −0.0176] *p*, = 3.58 × 10^−6^ and A4+LEARN (annualized *β* = −0.0205 PHC_MEM units/year, 95% CI [−0.0344, −0.0067], *p* = 0.0037), with slightly smaller effect estimates (Supplementary Figure S1). In A4+LEARN, the late-stage group also declined faster on PACC (annualized *β* = −0.1324 PACC units/year, 95% CI [−0.1900, −0.0749], *p* = 6.64 × 10^−6^; Figure 4E). Within this cohort, stratification was stronger for PACC than for PHC_MEM. Because all A4+LEARN participants were cognitively unimpaired at baseline, this likely reflects the greater sensitivity of PACC to subtle cognitive change in this population. Notably, DAPS was trained in ADNI, where participants span the cognitively normal to impaired range, and was then applied without retraining to A4+LEARN, a cohort composed entirely of cognitively unimpaired individuals. That the staging score retained clear stratification of subsequent decline under this shift supports its value for prognostic assessment in the earliest stage of Alzheimer’s disease.

**Figure 4.**
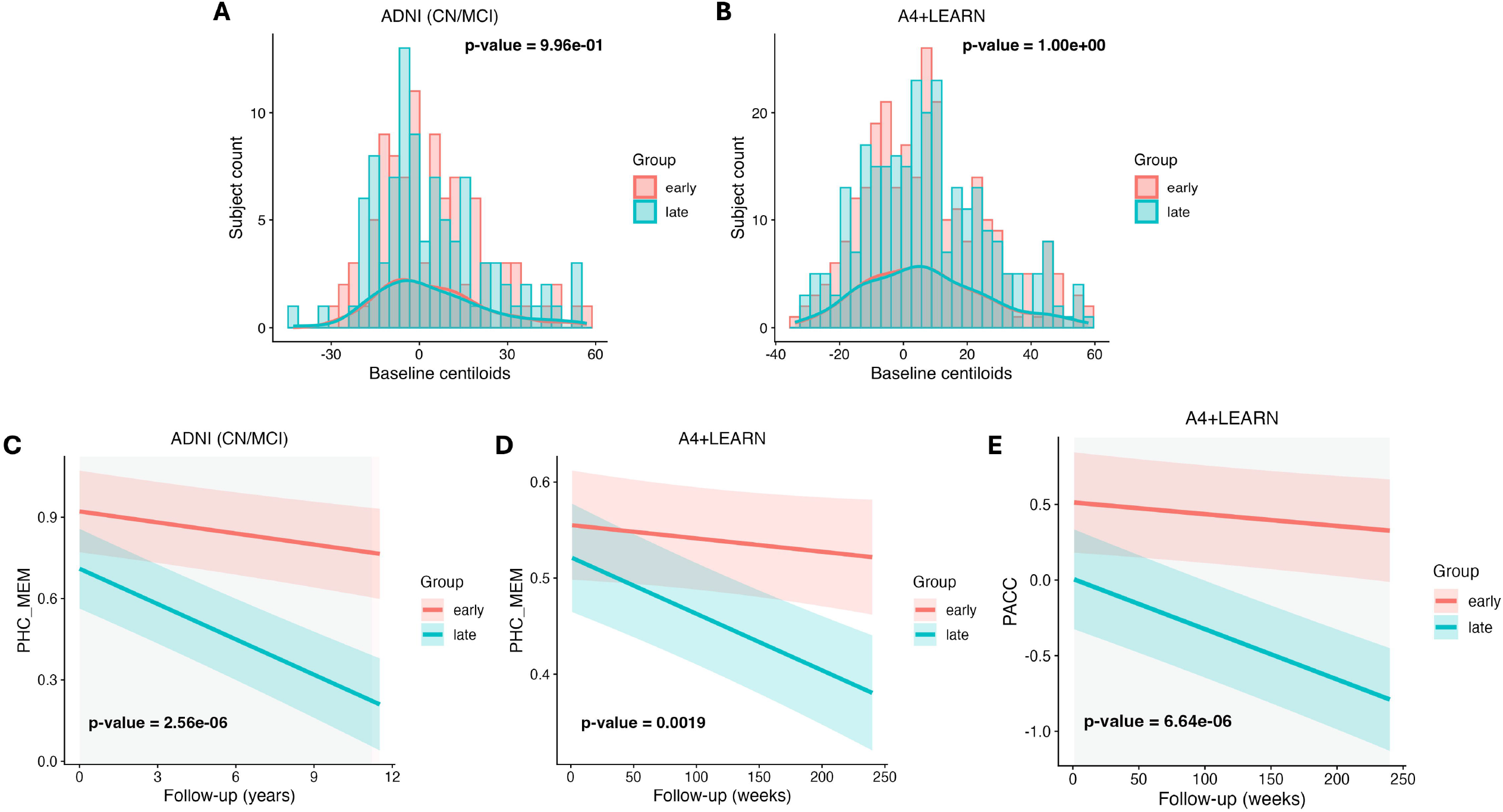
Longitudinal cognitive decline of early- and late-groups, stratified using the DAPS staging breakpoint (0.370). (A–B) In both ADNI test set (CN/MCI only) and A4+LEARN cohort, participants in early- and late-groups were sampled for matched baseline Centiloid distributions. *P* values were obtained from two-sample Kolmogorov–Smirnov tests. (C–D) Longitudinal cognitive trajectories (PHC_MEM) of early- and late-subgroups in ADNI test set (CN/MCI only) and A4+LEARN cohort. (E) Longitudinal cognitive trajectories (PACC) of early- and late-subgroups in A4+LEARN cohort. Across panels C–E, *P* values correspond to the group-by-follow-up-time interaction from the linear mixed-effects models.

## 4. Discussion

In this study, we developed DAPS, an end-to-end framework that directly estimates a continuous amyloid staging score from regional amyloid PET measurements. The proposed staging score demonstrated improved longitudinal consistency, exhibited a reproducible nonlinear relationship with global amyloid burden, and identified differences in subsequent cognitive decline among individuals with similar baseline global amyloid burden. Together, these findings support the potential use of continuous regional amyloid staging for characterizing the early phase of Alzheimer’s disease.

The relationship between the DAPS staging score and global amyloid burden provides insight into the biological interpretation of the proposed staging framework. During the very early stage of amyloid accumulation, amyloid deposition is typically confined to a small number of cortical regions, and the regional amyloid burden remains relatively low. Consequently, averaging regional measurements into a single global measure may reduce sensitivity to these localized changes. In contrast, the DAPS staging score is derived from the spatial pattern of regional amyloid PET measurements, thereby preserving regional information that may not yet substantially influence the global average. After the identified breakpoint, amyloid deposition becomes more widespread and regional amyloid burden increases across multiple cortical regions. At this stage, both the global amyloid burden and the DAPS staging score reflect the progressive accumulation of cortical amyloid and therefore increase in parallel. This interpretation is consistent with previous amyloid PET studies showing that amyloid accumulation initially occurs in a restricted set of cortical regions and that early regional abnormalities may remain undetected by conventional global amyloid PET classification^18,19^. The regional contribution analysis within early stage (i.e., small DAPS staging score) provides additional biological support for this interpretation. Regions with the highest contribution scores included the precuneus, posterior cingulate, inferior parietal cortex, medial orbitofrontal cortex, and cingulate regions. These regions largely overlap with those identified as early sites of amyloid accumulation in previous studies, including the precuneus, posterior cingulate, medial orbitofrontal, and parietal cortices^20^. Collectively, these findings suggest that the DAPS staging score may provide additional characterization of early amyloid accumulation beyond what is provided by global amyloid burden alone.

The ability to estimate a continuous staging score for previously unseen individuals extends the potential clinical application of DAPS beyond exploratory disease progression modeling. Continuous staging may be particularly valuable during the early phase of Alzheimer’s disease.

Recent studies suggest that spatial measures of amyloid deposition may improve prognostic assessment beyond conventional global amyloid measures^21^ Consistent with this emerging evidence, our results suggest that the baseline DAPS staging score provides complementary prognostic information beyond global amyloid burden alone. Such information may improve individual risk stratification and facilitate participant enrichment for observational studies and disease-modifying clinical trials targeting early Alzheimer’s disease. The consistency of these associations across both ADNI and the independent A4+LEARN cohort further supports the potential generalizability of the proposed staging framework.

Several limitations should be acknowledged. DAPS was developed and evaluated using processed and harmonized amyloid PET data provided through the ADSP Phenotype Harmonization Consortium. Although this harmonized framework enabled direct external validation across ADNI and A4+LEARN, application to other amyloid PET datasets would require comparable preprocessing and harmonization procedures. Extending the proposed framework to operate directly on raw amyloid PET images represents an important next step toward broader clinical applicability. Both cohorts were also imaged with florbetapir, which has a narrower dynamic range and shows reduced separation in the low and intermediate amyloid range {{Properzi, 2019 #22}}. DAPS cannot recover contrast that the tracer does not provide, although learning from regional amyloid patterns may partially offset the limited contrast of single regions. Future studies should further evaluate its generalizability across additional cohorts, amyloid tracers, and imaging protocols. Despite these limitations, the present study on regional measures already demonstrates potential in characterizing early-stage amyloid accumulation and in identifying individuals at different risks of future cognitive decline. These findings support further investigation of the DAPS score in observational studies and therapeutic trials targeting early Alzheimer’s disease.

## Acknowledgments

Data collection and sharing for the Alzheimer’s Disease Neuroimaging Initiative (ADNI) is funded by the National Institute on Aging (National Institutes of Health Grant U19AG024904). The grantee organization is the Northern California Institute for Research and Education. In the past, ADNI has also received funding from the National Institute of Biomedical Imaging and Bioengineering, the Canadian Institutes of Health Research, and private sector contributions through the Foundation for the National Institutes of Health (FNIH) including generous contributions from the following: AbbVie, Alzheimer’s Association; Alzheimer’s Drug Discovery Foundation; Araclon Biotech; BioClinica, Inc.; Biogen; BristolMyers Squibb Company; CereSpir, Inc.; Cogstate; Eisai Inc.; Elan Pharmaceuticals, Inc.; Eli Lilly and Company; EuroImmun; F. Hoffmann-La Roche Ltd and its affiliated company Genentech, Inc.; Fujirebio; GE Healthcare; IXICO Ltd.; Janssen Alzheimer Immunotherapy Research & Development, LLC.; Johnson & Johnson Pharmaceutical Research & Development LLC.; Lumosity; Lundbeck; Merck & Co., Inc.; Meso Scale Diagnostics, LLC.; NeuroRx Research; Neurotrack Technologies; Novartis Pharmaceuticals Corporation; Pfizer Inc.; Piramal Imaging; Servier; Takeda Pharmaceutical Company; and Transition Therapeutics.

The ADSP Phenotype Harmonization Consortium (ADSP-PHC) is funded by NIA (U24 AG074855, U01 AG068057 and R01 AG059716). The Anti-Amyloid Treatment in Asymptomatic Alzheimer’s study (A4 Study), a secondary prevention trial in preclinical Alzheimer’s disease, aiming to slow cognitive decline associated with brain amyloid accumulation in clinically normal older individuals. The A4 Study is funded by a public-private-philanthropic partnership, including funding from the National Institutes of Health-National Institute on Aging, Eli Lilly and Company, Alzheimer’s Association, Accelerating Medicines Partnership, GHR Foundation, an anonymous foundation and additional private donors, with in-kind support from Avid and Cogstate. The companion observational Longitudinal Evaluation of Amyloid Risk and Neurodegeneration (LEARN) Study is funded by the Alzheimer’s Association and GHR Foundation. The A4 and A4-LEARN Studies are led by Dr. Reisa Sperling at Brigham and Women’s Hospital, Harvard Medical School and Dr. Paul Aisen at the Alzheimer’s Therapeutic Research Institute (ATRI), University of Southern California. The A4 and LEARN Studies are coordinated by ATRI at the University of Southern California, and the data are made available through the Laboratory for Neuro Imaging at the University of Southern California. The participants screening for the A4 Study provided permission to share their de-identified data in order to advance the quest to find a successful treatment for Alzheimer’s disease. We would like to acknowledge the dedication of all the participants, the site personnel, and all of the partnership team members who continue to make the A4 and LEARN Studies possible. The complete A4 Study Team list is available on: a4study.org/a4-study-team.

## Consent Statement

Data used in this study were obtained from the Alzheimer’s Disease Neuroimaging Initiative (ADNI) and ADSP phenotype harmonization cohort. All participants provided written informed consent at enrollment, and institutional review board approval was obtained at each participating site. The present study is a secondary analysis of de-identified data.

## Data Availability

Harmonized amyloid PET data analyzed in this study were obtained respectively from the Alzheimer’s Disease Neuroimaging Initiative (ADNI) data portal (https://ida.loni.usc.edu) and through the Alzheimer’s Disease Sequencing Project (ADSP) Phenotype Harmonization Consortium. Access to these datasets is subject to the approval by the corresponding data repositories.

## Code Availability

Pre-trained model together with best parameters is shared through GitHub (https://github.com/StochasticAgCl/DAPS).

