## Supplemental Figure S1 for "Continuous amyloid staging complements global amyloid burden in early Alzheimer’s disease"

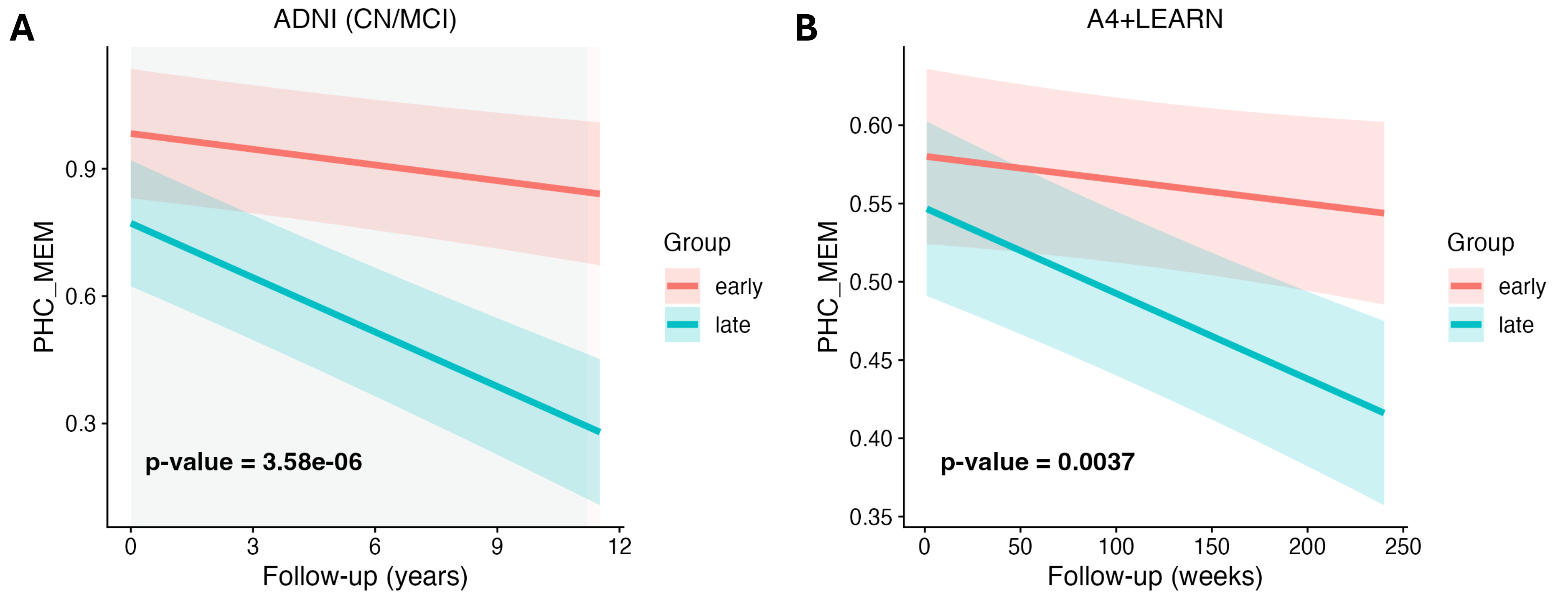


**Figure S1.** Longitudinal cognitive trajectories (PHC_MEM) in ADNI test set and independent A4+LEARN cohort, respectively, were estimated using linear mixed-effects models without considering measurement precision (i.e., standard error of PHC_MEM). The same matched participants and observations as in the corresponding precision-aware analyses were included. In both cohorts, PHC_MEM trajectories differed significantly between the early- and late-stage groups. *P* values correspond to the group-by-follow-up-time interaction from the linear mixed-effects models.
