## Supplementary material for "Continuous amyloid staging complements global amyloid burden in early Alzheimer’s disease": Table 1

Table 1: Demographic information of all participants.

| ADNI | | | | |  | A4+LEARN |
| --- | --- | --- | --- | --- | --- | --- |
| Baseline Diagnosis | CN | MCI | AD | Unknown |  | All |
| Number | 409 | 531 | 188 | 33 |  | 3414 |
| Gender (Male/Female) | 191/218 | 313/218 | 111/77 | 16/17 |  | 2043/1371 |
| Baseline Age (mean ± sd) | 74.15 ± 6.77 | 73.00 ± 7.84 | 75.48 ± 7.67 | 74.55 ± 8.86 |  | 71.21 ± 4.70 |
| Years of Education (mean ± sd) | 16.67 ± 2.56 | 16.11 ± 2.72 | 15.85 ± 2.62 | 15.82 ± 2.69 |  | 16.65 ± 2.85 |
